# Cropping systems impact changes in soil fungal, but not prokaryote, alpha-diversity and community composition stability over a growing season in a long-term field trial

**DOI:** 10.1101/2020.03.15.992560

**Authors:** D.R. Finn, S. Lee, A. Lanzén, M. Bertrand, G.W. Nicol, C. Hazard

## Abstract

Crop harvest followed by a fallow period can act as a disturbance on soil microbial communities. Cropping systems intended to improve alpha-diversity of communities may also confer increased compositional stability during succeeding growing seasons. Over a single growing season in a long-term (18 year) agricultural field experiment incorporating conventional (CON), conservation (CA), organic (ORG) and integrated (INT) cropping systems, temporal changes in prokaryote, fungal and arbuscular mycorrhizal fungi (AMF) communities were investigated overwinter, during crop growth and at harvest. While certain prokaryote phyla were influenced by cropping system (*e.g.* Acidobacteria), the community as a whole was primarily driven by temporal changes over the growing season as distinct overwinter and crop-associated communities, with the same trend observed regardless of cropping system. Species-rich prokaryote communities were most stable over the growing season. Cropping system exerted a greater effect on fungal communities, with alpha-diversity highest and temporal changes most stable under CA. CON was particularly detrimental for alpha-diversity in AMF communities, with AMF alpha-diversity and stability improved under all other cropping systems. Practices that promoted alpha-diversity tended to also increase the similarity and temporal stability of soil fungal (and AMF) communities during a growing season, while prokaryote communities were largely insensitive to management.

## 1. Introduction

The soil microbial community is primarily controlled by both plant-dependent factors and soil physico-chemistry (Philippot *et al*., 2013), particularly physico-chemistry related to soil fertility management (Guo *et al*., 2020, Semenov *et al*., 2020). It is a complex assembly including Bacteria, Archaea, obligate plant-symbiont arbuscular mycorrhizal fungi (AMF) and saprotrophic, non-mycorrhizal fungi. Bacteria regulate many essential processes, including nitrogen fixation (Rosswall, 1982), nitrification (Kowalchuk & Stephen, 2001), decomposition of complex organic matter (*i.e.* cellulose, hemicellulose, chitin) (Schimel & Schaeffer, 2012) and emissions of greenhouse gases methane and nitrous oxide (Hanson & Hanson, 1996, Barnard *et al*., 2005). In the soil environment, methanogenic archaea produce methane and members of the phylum Thaumarchaeota oxidise ammonia (Prosser & Nicol, 2008) with their activity often distinct from bacterial ammonia oxidisers. AMF are vital for the growth of many plant species by facilitating nutrient uptake (*e.g.* phosphorus and nitrogen) from soil and transfer to roots (Koide & Kabir, 2000), and also improve soil aggregation and carbon sequestration via hyphal growth (Borie *et al*., 2006). Finally, saprotrophic fungi also assist in the decomposition and recycling of complex organic matter (Setala & McLean, 2004). The combined activity of these microorganisms benefit plant-growth, often in exchange for root exudates as an organic carbon source (Cordovez *et al*., 2019).

Relatively high alpha-diversity (*e.g.* richness, evenness) in soil microbial communities is associated with improved plant net primary production, nitrogen mineralisation and microbial biomass production, amongst other functions (Delgado-Baquerizo *et al*., 2016). Therefore, cropping systems that have the potential to improve alpha-diversity are desirable. Tillage, for example, has a particularly detrimental effect on AMF community richness (Sale *et al*., 2015, Banerjee *et al*., 2019). Mineral nitrogen fertilisation favours Actinobacteria, Proteobacteria and ammonia oxidising bacteria (AOB) over ammonia oxidising archaea (AOA) (Fierer *et al*., 2012, Illescas *et al*., 2020). Organic fertilisation increases richness of Verrucomicrobia, Planctomycetes, Acidobacteria and if manure is applied, Firmicutes associated with the gut (Wessen *et al*., 2010). Organic amendments can alleviate negative impacts of salt-stress on alpha-diversity (Szoboszlay *et al*., 2019). Crop rotations have the potential to increase bacterial alpha-diversity, while simultaneously decreasing that of fungi (Zhou *et al*., 2017). Pesticides have the potential to decrease nitrogen-fixing and nitrifying bacteria, among others (Johnsen *et al*., 2001). Permanent cover crops structure the community in many ways: a) their root systems change soil structure that stabilises soil moisture content and increases overall microbial richness (Vukicevich *et al*., 2016); b) AMF-associated with cover crops remain stable in the soil, and can even transfer nutrients to commercial crops when they are present (Cheng & Baumgartner, 2006); and c) the rhizosphere of cover crops can promote richness of specific root disease-suppressive pseudomonads and fungal *Trichoderma* spp. (Vukicevich *et al*., 2016). Employing cropping systems that utilise management practices aimed at maximising species richness can thus maximise the benefits of soil microbial function on plant growth.

A common practice in agricultural systems that cultivate annual crops sown and harvested within a year, *e.g.* wheat (*Triticum aestivum* L.) or rapeseed (*Brassica napus*), is to have a fallow period between harvest and sowing of the next season’s crop. A crop is not grown within this fallow period. Due to the importance of plant-microbe interactions in shaping the soil microbial community, the removal of an active root system at harvest and subsequent fallow period likely acts as a major disturbance event. Where cropping systems that foster higher alpha-diversity are employed, the negative effect of disturbance on community composition may be less pronounced, *i.e.* they would be relatively more stable. This is an example of the insurance hypothesis, whereby communities that display greater species richness should be more robust to disturbance (Ives *et al*., 2000) and ultimately ensure more stable functionality. Similarly, low agriculture intensity would be expected to promote relatively higher alpha-diversity, and to yield more stable relationships between individual taxa (i.e. network connections; e.g. Banerjee *et al*., 2019).

The La Cage long-term experimental agricultural site was established by the Institut National de la Recherche pour l’Agriculture, l’Alimentation et l’Environnement (INRAE) in 1998 in Versailles, France. Its purpose is to compare the performance of four cropping systems; conventional (CON) *versus* those with a lesser intensity of disturbance - integrated (INT), conservation (CA) and organic (ORG). CA has been shown to increase soil organic carbon stocks, without increasing soil respiration rate, relative to other cropping systems (Autret *et al*., 2016). Additionally, CA improved bacterial, fungal, earthworm and arthropod biomass relative to CON, while only ORG increased bacterial biomass relative to other cropping systems (Henneron *et al*., 2015). The benefits of CA are thought to be primarily derived from the incorporation of a permanent cover crop, which is unique to the CA cropping system and absent from CON, INT and ORG. The La Cage site therefore represents a valuable long-term model system to compare how various agricultural practices affect the alpha-diversity of microbial communities and how they change over a growing season. To address this, microbial community composition analyses were performed for prokaryotes, fungi and specifically for AMF on CON, CA, INT and ORG soils overwinter (December), part-way through the growing season (May) and immediately prior to harvest (July). As the CON, CA, INT and ORG soil communities have been under the dual selection pressures of their respective cropping system and cyclical annual harvesting for 18 years, it was expected that both factors would play a role in determining composition, and the development of potential stable relationships between taxa.

We first sought to investigate whether cropping systems have differentially affected the alpha-diversity (as richness and Shannon index) of soil microbial communities in this long-term experimental system. Secondly, we sought to determine whether stable relationships between individual taxa have emerged under different cropping systems. Finally, we investigated whether cropping systems that promoted alpha-diversity would also promote relatively more stable microbial community compositions over time. This stability was measured as compositional similarity, whereby communities that remained similar over time were considered to be relatively stable, while those that demonstrated greater dissimilarity were considered less stable.

## 2. Materials and methods

### 2.1. La Cage experimental site

A detailed presentation of the La Cage long term trial has been reported previously (Debaeke *et al*., 2009). A diagram of the La Cage experimental design is shown in Figure 1. The site is located 15 km southwest of Paris (48◦48’ N, 2◦08’ E). The soil is a deep loamy, well-drained Luvisol (WRB, 2015). Texture includes: 15 – 18.4% clay (< 2 μm particle size diameter), 16.5 – 20.2% fine silt (2 – 20 μm), 30.3 – 43.2% coarse silt (20 – 50 μm) and 18.4 – 31.2% sand (50 – 200 μm). Table 1 outlines specific practices, crop production (wheat, pea and rapeseed t ha^-1^ yr^-1^) and soil properties (total organic carbon TOC, total nitrogen TN, C:N ratio, pH and cation exchange capacity CEC) that differ between the four cropping systems. Briefly, CON is characterised by frequent tillage, pesticide application and high level of mineral N fertilisation. CA is no-till, infrequent pesticide application, has a permanent cover crop and receives N from mineral sources and a legume rotation. INT involves reduced tillage, infrequent pesticide application, and receives a reduced level of mineral N fertilisation. Finally, ORG involves frequent tillage, no pesticide application, and receives N from a legume rotation. Soil physico-chemical analyses were measured by INRAE in 2014, independent of this study, as described (Debaeke *et al*., 2009) from two pseudo-replications from subplots of each cropping system, from two independent blocks (*n* = 4 per cropping system). Annual crops wheat (*Triticum aestivum* L.) and rapeseed (*Brassica napus*) were grown on independent subplots (0.56 ha) within each block and undergo annual rotations between subplots at La Cage. Rapeseed was sown in August, and wheat in October. The four cropping systems (CON, CA, INT and ORG) were applied to each of the wheat/rapeseed subplots. The cover crops under CA were red fescue (*Festuca rubra*) and alfalfa (*Medicago sativa*). The cover crop is permanently maintained, but mechanically or chemically supressed prior to sowing wheat and rapeseed (i.e. a direct seeding living mulch-based system). For CON, INT and CA, mineral N application occurred yearly, with the exception of a rotating subplot (12 m x 20 m) that did not receive mineral N that year. Mineral fertiliser was applied in February. The ORG system does not involve fertilisation from mineral N sources or manure application, rather N comes from a legume rotation. During this study, ORG had an alfalfa crop rotation (sown in August) in place of rapeseed. Crops were harvested in July.

**Figure 1:**
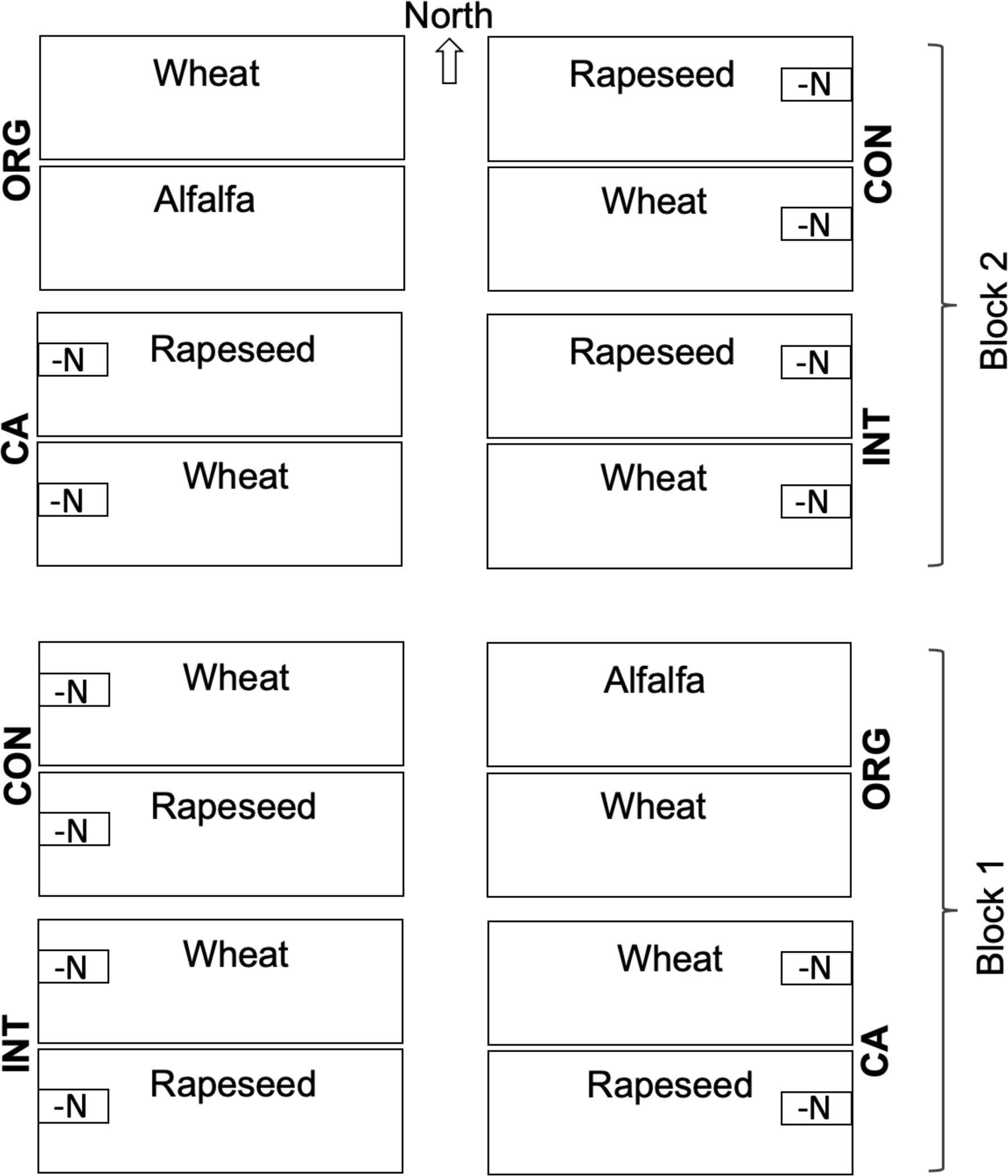
Diagram of the La Cage experimental design. Two independent blocks (1 and 2) were sampled. Each block consists of four plots (1.12 ha each) in which the four cropping systems (Conventional, CON, Conservation, CA, Integrated, INT, and Organic, ORG) were randomly applied. Each plot is split into two subplots (0.56 ha) with annual rotations of wheat and rapeseed. CA and ORG also have legume rotations. During this study, ORG had an alfalfa crop rotation. For CON, INT and CA, mineral N application occurred yearly, with the exception of a rotating subplot (12 m x 20 m) that does not receive mineral N (-N) that year (i.e. applied in 2015 *versus* applied in 2016). Five pseudo-replicate samples were taken from each crop (wheat/rapeseed) and N (2015/2016) subplot, for CON, CA and INT (*n* = 20 each) from each block. Ten pseudo-replicate samples were taken from each crop (wheat/alfalfa) subplot for ORG (*n* = 10) from each block.

**Table 1:** Management practices, crop production and soil properties of the La Cage cropping systems. Superscript letters indicate Tukey’s HSD outcomes between cropping systems at the p < 0.05 level.

|  | CON | CA | INT | ORG |
| --- | --- | --- | --- | --- |
| Tillage<sup>1</sup> | Annual | None | Biannual | Annual |
| N fertilisation | Mineral-high | Mineral-reduced | Mineral-reduced | None |
| Pesticides | Frequent | When nec. | When nec. | None |
| Cover crop | No | Yes | No | No |
| Legume rotation | No | Yes | No | Yes |
| Wheat (t ha<sup>-1</sup> yr<sup>-1</sup>) | 9.7 | 6.7 | 8.9 | 5.4 |
| Pea (t ha<sup>-1</sup> yr<sup>-1</sup>) | 4.2 | 3.7 | 4.5 | 2.6 |
| Rapeseed (t ha<sup>-1</sup> yr<sup>-1</sup>) | 4.5 | - | 3.8 | 0.8 |
| TOC | 11.1 ± 0.9<sup>a,b,c,d</sup> | 13.2 ± 2.8<sup>a,b,c</sup> | 11 ± 1.6<sup>a,b,c,d</sup> | 10.4 ± 0.5<sup>a,b,d</sup> |
| TN | 0.9 ± 0.1<sup>a</sup> | 1.1 ± 0.2<sup>b</sup> | 0.9 ± 0.1<sup>a</sup> | 0.9 ± 0.1<sup>a</sup> |
| C:N ratio | 12.4 ± 1<sup>a</sup> | 11.5 ± 0.1<sup>a</sup> | 11.8 ± 0.2<sup>a</sup> | 11.6 ± 0.3<sup>a</sup> |
| pH | 7.3 ± 0.3<sup>a</sup> | 6.9 ± 0.2<sup>a</sup> | 7.1 ± 0.4<sup>a</sup> | 7 ± 0.2<sup>a</sup> |
| CEC | 9.9 ± 1<sup>a</sup> | 9.6 ± 0.3<sup>a</sup> | 9.5 ± 1<sup>a</sup> | 9.2 ± 0.3<sup>a</sup> |

Soil properties were measured in all subplots in 2014 at INRAE.
<sup>1</sup>Tillage occurs soon after crop harvest.

### 2.2. Soil sampling

From the CON, CA and INT cropping systems, five pseudo-replicate soil samples were taken from each subplot, respectively, from each combination of crop (wheat *versus* rapeseed (or alfalfa for ORG)) and mineral N rotation subplot (applied in 2015 *versus* applied in 2016). This yielded 20 pseudo-replications per CON, CA and INT cropping system. As ORG did not involve a mineral N rotating subplot, only 10 pseudo-replicate samples (five from wheat, five from alfalfa) were collected. Soil samples, at a diameter and depth of 4 and 20 cm, were randomly collected between densely distributed plants, with plot edges avoided and at least 5 meters between samples. Soil samples were collected in December 2015 (overwinter), May 2016 (during growing season) and July 2016 (at harvest). Thus, a total of 70 samples were collected at each of the three time points, for a total of 210 samples. Finally, two spatially independent blocks were sampled. These independent blocks represented true replications of the cropping systems, as opposed to the pseudo-replicate samples taken within subplots, within each block. In total, 420 soil samples were collected for molecular analyses. Soil samples were sieved (3.35 mm) and stored at -20°C.

### 2.3. Molecular analyses

DNA was extracted from 0.5 g soil using the MoBio PowerSoil DNA Kit (MoBio, Carlsbad, CA, USA) following the manufacturer’s protocol. Prokaryote- and AMF-specific amplification of fragments of the small subunit rRNA gene (16S and 18S respectively) was carried out using the primer pairs 515F (Parada *et al*., 2016)/806R (Apprill *et al*., 2015) and AMV4.5NF/AMDGR (Sato *et al*., 2005), respectively. Amplification of the fungal internal transcribed spacer 1 (ITS1) region was carried out with the primer pair ITS1F (Gardes & Bruns, 1993)/ITS2 (White *et al*., 1990). Primers had Illumina adaptor sequences attached. PCR was performed in 27 μl reactions with 22.5 μl of Invitrogen Platinum PCR SuperMix (Thermo Fisher, Carlsbad, CA, USA), 1.0 μl of forward and reverse primers (10 μM), and 2.5 μl of template DNA (5 ng/μl) on a Biometra T1 thermocycler (Biomentra GmbH, Göttingen, Germany). Thermocycling conditions for 16S rRNA gene and ITS assays were as follows: 94°C for 3 min; 35 cycles at 94°C for 45 s, 50°C for 1 min, 72°C for 90 s; and 72°C for 10 min. For 18S rRNA genes, 95°C for 5 min; 35 cycles at 95°C for 45 s, 56°C for 45 s, 72°C for 1 min; and 72°C for 7 min was used. Amplicons were bead purified using Agencourt AMPure XP (Beckman Coulter, Villepinte, France), followed by indexing PCR using the Nextera XT Index Kit (Illumina, San Diego, CA, USA) following manufacture’s recommendations. Indexed amplicons were bead purified using Agencourt AMPure XP and quantified using a μDrop Plate (Thermo Fisher, Carlsbad, CA, USA). Equimolar concentrations of samples were pooled and sequenced on an Illumina MiSeq sequencer with V2 2x150 bp paired-end chemistry. The raw sequences, with associated metadata, are available at NCBI BioProject under accession PRJN609408.

### 2.4. Bioinformatic analyses

Sequencing data was manually inspected using FastQC (Andrews, 2010), and processed using a workflow previously described in Martínez-Santos *et al*. (2018), modified for fungal ITS and AMF 18S rRNA gene analyses as described in detail below. Sequence read pairs were first merged (95% of read pairs merged), using *vsearch* allowing for a maximum of 20 differences and “staggered” read overlaps (*- fastq_maxdiffs 20 -fastq_allowmergestagger*) (Rognes *et al*., 2016). Thereafter, paired reads were trimmed to remove forward and reverse primers using *cutadapt* (Martin, 2011). Sequences lacking the correct primer sequences were discarded (using the *- discard-untrimmed* argument of cutadapt). Paired 16S rRNA gene sequences were then trimmed to a length of 252 and shorter sequences were discarded using vsearch (*-fastq_filter -fastq_trunclen 252*). For fungal ITS and AMF 18S rRNA gene sequences, with a more variable amplicon length, no trimming was carried out but sequences shorter than 100 or 200 bp, respectively, were discarded (0.43% of merged reads). Further, for all amplicons, sequences with more than one expected error were discarded, using *vsearch* (*-fastq_maxee 1*). Sequence filtering as described above resulted in 87% (10,201,515 sequences), 90% (8,788,547 sequences) and 92% (5,463,549 sequences) of the raw 16S rRNA, ITS and 18S rRNA read pairs, respectively. Clustering and denoising was performed using Swarm v2 with default parameters and fastidious mode (*-f*) (Mahe *et al*., 2015), followed by reference based and *de novo* chimera checking using *vsearch* (UCHIME algorithm; Edgar *et al*., 2011), with the RDP Gold reference sequences for 16S rRNA, SilvaMod v106 for 18S rRNA (Lanzen *et al*., 2012) and UNITE (Koljalg *et al*., 2005) for ITS. Of the filtered reads, 7.5%, 0.3% and 0.3% for 16S rRNA, ITS and 18S rRNA, respectively, were chimeric. SWARM OTUs were then further subjected to clustering using *vsearch* with a cut-off of 97% minimum similarity, and singletons retained (0.13%, 0.01% and 0.01% of 16S rRNA, ITS and 18S rRNA reads, respectively). Taxonomic classification of representative OTU sequences was then carried out using CREST (Lanzen *et al*., 2012) with the Silva v123 reference database (Quast *et al*., 2013), except for ITS for which the UNITE database was used (Koljalg *et al*., 2005). OTUs not identified as bacteria or archaea, fungi and Glomeromycota were removed from the Prokaryote 16S rRNA (0.2% of reads), fungal ITS (10% of reads) and AMF 18S rRNA (70% of reads) datasets, respectively. The MaarjAM database was used for AMF taxa identification of the OTUs to genera (similarity ≥ 97%, query coverage ≥ 90 %, E-value ≤ 1e^-100^) (Opik *et al*., 2010).

### 2.5. Statistical analyses

All statistics were performed in R v4.0.0 (Team, 2013). Initially Tukey’s Honest Significant Difference (HSD) test was performed to test for differences in soil TOC, TN, C:N ratio, pH and CEC across cropping systems. Two pseudo-replicates from each subplot, per block, for each cropping system (*n* = 4 each) were tested.

Samples with less than 4000 prokaryote or fungal sequences, and less than 200 AMF sequences, were removed from the datasets prior to the following analyses (1, 4 and 19 samples were removed, respectively). Heatmaps showing the relative abundance (%) of phyla within prokaryotes, fungi and AMF, for each sample across time, cropping system and blocks (*n* = 420) were generated with the package ‘gplots’ (Warnes *et al*., 2019). Due to their extensive taxonomic diversity, Proteobacteria were displayed at the class level. Only taxonomic groups that comprised more than at least 5% of the total community are shown. Effects of time, cropping system, crop (wheat/rapeseed (alfalfa for ORG)), year of N application (2015/2016) in subplots and blocks (1/2) on compositional abundances at the Phylum level were performed with the ANOVA-like differential expression (ALDEx2) (Fernandes *et al*., 2013, Fernandes *et al*., 2014, Gloor *et al*., 2017) in R. Phyla abundances were centred log-ratio (CLR) transformed with 128 Monte-Carlo samplings and, when found to have non-normal CLR-based univariate distributions (Korkmaz *et al*., 2014), effects were tested with Kruskal-Wallis in ALDEx2 with the ‘aldex.kw’ function. Probability values were Benjamini-Hochberg adjusted to account for false discovery rate.

Species richness and Shannon indices presented were calculated using non-rarefied data (McMurdie and Holmes, 2014). Rarefying had no effect on resulting trends. Indices were visualised with box and whisker plots across time and between cropping systems. Linear mixed effects models were performed to test for differences in richness and Shannon index across time and between cropping system. Random effects were considered as crop, year of N application and block. Models were applied with the ‘lme4’ package as described (Bates *et al*., 2015). Principle components analysis (PCA) was performed on CLR-transformed OTU matrices based on Euclidean distance (Vermeesch *et al*., 2016) via the ‘vegan’ package (Oksanen *et al*., 2013). Zero handling of sparse matrices was corrected by adding 1 to all cells. Sample sizes were as above, with time, cropping system, crop, year of N application and block tested on CLR-transformed OTUs with PERMANOVA separately, with 999 permutations and based on Euclidean distance. Multivariate homogeneity of variance was tested as described (Korkmaz *et al*., 2014).

For network analyses, prokaryote, fungi and AMF OTUs were subsetted based on individual management practice to compare the occurrence and stability of significant taxon-taxon relationships over time (*n* = 120, 120, 120, 60 for CON, CA, INT and ORG, respectively). That is, each network based on a single cropping system included each of the three time points. Weighted networks were visualised from Spearman covariance matrices, with non-significant edge weights below the 99^th^ quantile trimmed, performed with the ‘igraph’ package (Csardi & Nepusz, 2006). Spearman correlation has been shown to yield robust networks, outperforming compositional-based correlation methods, particularly in sparsely connected networks (Hirano & Takemoto, 2019). Due to their relatively low number of nodes and possible edges, AMF were trimmed at the 95^th^ quantile. Nodes that lacked at least one edge were filtered out. Weighted networks were visualised with the force-directed Kamada-Kawai algorithm. Total nodes, edges (*i.e.* significant relationships), edge betweenness scores, clustering coefficients and diameter of each network was calculated as described (Csardi & Nepusz, 2006).

Jaccard Similarity (JS) was calculated for each sample over time, as a pairwise measure of community similarity between harvest and overwinter (July *versus* December), and harvest and during growing season (July *versus* May) communities. The JS was chosen as it is a robust and simple measure of pairwise similarity commonly employed in the comparison of ecological communities (Scheiner, 1992). By measuring the JS after disturbance (December) and part-way through the growing season (May) we sought whether communities under certain practices, such as CA, would be more similar to the harvest community than other practices, such as CON. Where communities displayed consistently high similarity, these were considered relatively stable, while those with consistently high dissimilarity were considered unstable.

Jaccard Similarity (JS) was calculated as:

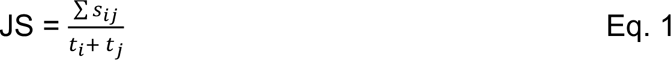

where *s* was the sum of shared OTUs in samples *i* and *j*, while *ti* and *tj* were the total OTUs in samples *i* and *j*, respectively. JS was calculated at the genus level for prokaryotes and fungi, and at the individual OTU level for AMF due to their relatively low richness. The July (harvest) time point for each sample was considered *i*, while *j* iterated through the December (overwinter) and May (part-way through cropping) time points. The JS of wheat *versus* rapeseed/alfalfa crop or year of N application were not compared, as each sample was only compared to its corresponding cropping system/crop/year of N application sample over time. Linear mixed effects models were performed to confirm whether cropping system affected JS at December and May time points, with crop, year of N application and block considered as random effects.

## 3. Results

### 3.1. Soil properties and microbial community composition

Table 1 lists soil properties that differed among the four cropping systems. TOC was greatest under CA and lowest under ORG. Only CA had significantly greater TN than the other schemes. The C:N ratio, soil pH and CEC did not differ based on management. Properties did not differ between blocks (data not shown).

Time was considered as a categorical variable with three levels: December sampling during winter with sown crops, May sampling part-way through the growing season and July sampling just prior to crop harvest. Relative abundances of dominant taxonomic groups are shown in Figure 2. The prokaryotic groups with relative abundance greater than 5% were Acidobacteria, Actinobacteria, Alphaproteobacteria, Bacteroidetes, Betaproteobacteria, Gammaproteobacteria, Deltaproteobacteria, Firmicutes, Planctomycetes, Thaumarchaeota and Verrucomicrobia. These relatively abundant groups were not affected by crop, N application or block (Kruskal-Wallis *p* > 0.05). Both time and cropping system had effects on the relative abundance of most prokaryote phyla, with time generally conveying a stronger effect. Cropping system affected the relative abundances of the dominant fungal Ascomycota, while most other phyla were primarily affected by time. Fungi were not affected by crop, N application or block (Kruskal-Wallis *p* > 0.05). Glomerales and Diversisporales were the most dominant AMF. Both were affected by time, while cropping system had no effect on AMF at the Order level. The Diversisporales and Glomerales were affected by crop (Kruskal-Wallis *p* < 0.05) and Diversisporales by block (Kruskal-Wallis *p* = 0.001).

**Figure 2:**
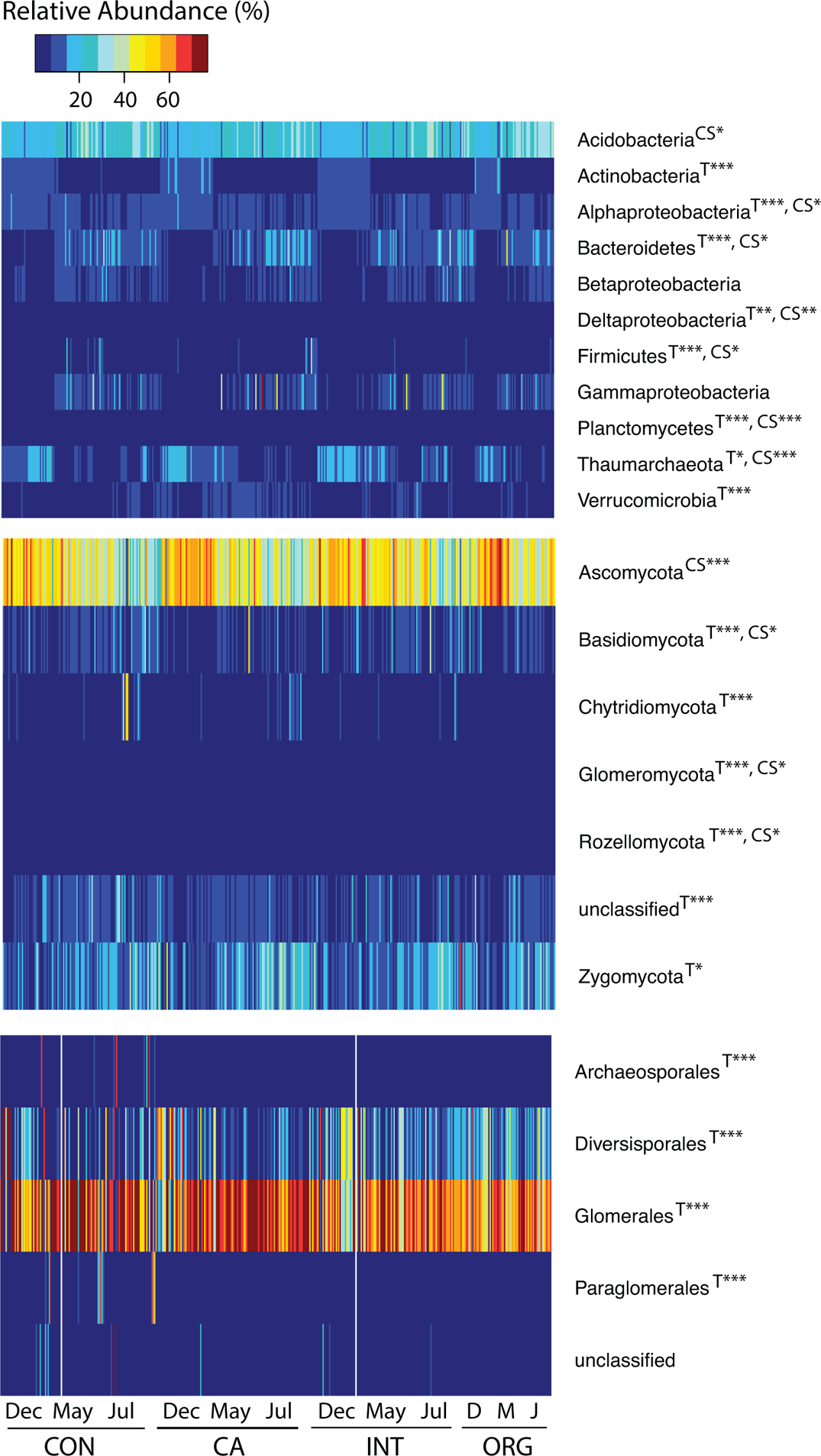
Heatmap comparing relative abundances of prokaryote and fungal phyla, and AMF orders, across time and between cropping system. A total of 420 samples are shown. Units are relative abundance (%) per sample. Relative abundance was calculated separately for prokaryote, fungal and AMF communities. Where taxa differed over time (T) or by cropping system (CS) Kruskal-Wallis tests on centred log-ratio transformed relative abundances are shown as: *p* < 0.05 (*); *p* = 0.001 (**); *p* < 0.001 (***).

Prokaryote richness and Shannon indices followed the trend Dec < May < July regardless of cropping system (Figure 3a). Both measurements increased in May and July relative to December (*p* < 0.05, Table 2) while cropping system had no effect on prokaryote alpha-diversity. Only year of N application had an appreciable random effect on prokaryote alpha-diversity (8.4 and 6.3% for richness and Shannon, respectively). Fungal richness was significantly highest in July, and interestingly, decreased in May relative to December (Table 2). The CA and INT systems had positive effects on fungal richness. The fungal Shannon index indicated negative effects of ORG and time between December and May. Random effects were very marginal for fungal diversity (< 2.4% for each). The AMF richness and Shannon were most sensitive to time and cropping system (Table 2). Relative to CON, the CA, INT and ORG systems all had strong positive effects on AMF alpha-diversity which continuously increased over time. However, crop (wheat *versus* rapeseed (alfalfa for ORG)) played an important background role for AMF, accounting for 15.4 – 22.8% of observed variation in diversity as a random effect.

**Figure 3:**
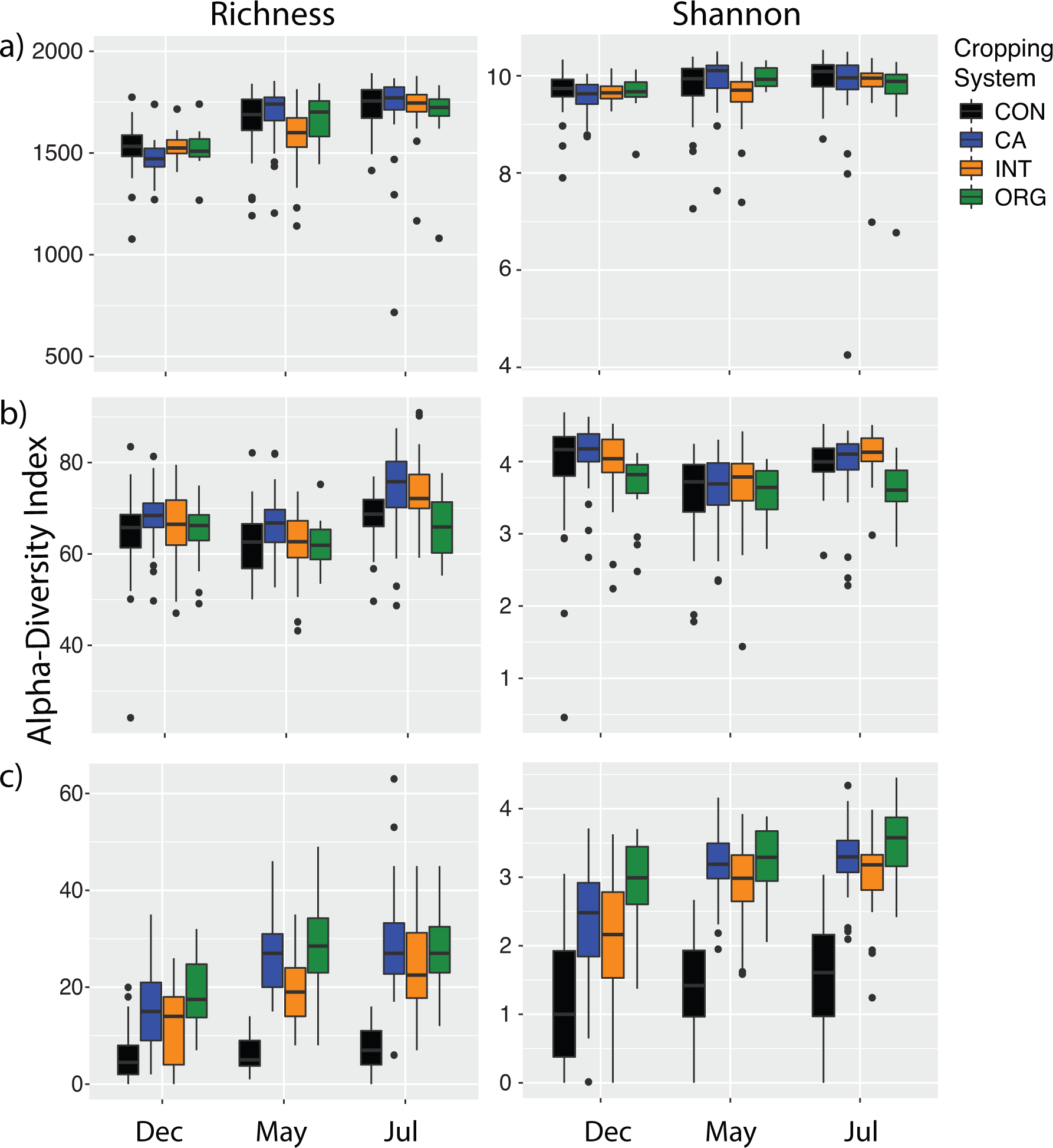
Observed OTU richness and Shannon indices for: a) prokaryotes, b) fungi and c) AMF.

**Table 2:** Mixed effects models of cropping system effects on prokaryote, fungal and AMF alpha-diversity indices. Also shown are the intercepts for each treatment and time point, with statistical significance noted as: (***) *p* < 0.001, (**) *p* = 0.001, (*) *p* < 0.05. Variance explained by random effects crop, year of N application and block are also shown.

|  |  |  |  |  |  | Cropping System |  |  |  | Time |  |  | Variance explained by random effects (%) |  |  |
| --- | --- | --- | --- | --- | --- | --- | --- | --- | --- | --- | --- | --- | --- | --- | --- |
|  | Intercept | Std.Error | t value | p value |  | CON | CA | INT | ORG | Dec | May | Jul | Crop | Nitrogen | Block |
| Prokaryotes |  |  |  |  |  |  |  |  |  |  |  |  |  |  |  |
| Richness | 1514.925 | 32.22 | 47.02 | 0.001** |  | 1514.925 | 1509.135 | 1491.507 | 1483.173 | 1514.925 | 1657.867*** | 1731.155*** | 0.8 | 8.39 | 0.003 |
| Shannon | 9.67 | 0.13 | 76.53 | < 0.001*** |  | 9.67 | 9.62 | 9.572 | 9.551 | 9.67 | 9.835* | 9.868** | 1.23 | 6.3 | 0.28 |
| Fungi |  |  |  |  |  |  |  |  |  |  |  |  |  |  |  |
| Richness | 63.98 | 1.32 | 48.44 | < 0.001*** |  | 63.98 | 68.68*** | 66.28* | 64.14 | 63.98 | 61.37** | 69.07*** | 0.37 | 2.32 | 1.44 |
| Shannon | 3.91 | 0.06 | 11.8 | < 0.001*** |  | 3.9 | 4 | 4.01 | 3.73* | 3.9 | 3.56*** | 3.902 | 0 | 0 | 0.39 |
| AMF |  |  |  |  |  |  |  |  |  |  |  |  |  |  |  |
| Richness | 0 | 2.8 | 0.35 | 0.8 |  | 0 | 17.22*** | 12*** | 21.38*** | 0 | 6.85*** | 8.89*** | 22.83 | 1.3 | 0 |
| Shannon | 0.8 | 0.21 | 3.91 | 0.03* |  | 0.8 | 2.35*** | 2.09*** | 2.91*** | 0.8 | 1.41*** | 1.56*** | 15.42 | 0 | 1.83 |

Figure 4 displays the PCA of prokaryote (a), fungal (b) and AMF (c) community compositions. Prokaryotes were primarily affected by time (PERMANOVA, R^2^ = 0.13, *p* < 0.001) as demonstrated by distinct separation of all December communities (circles, Figure 4a). Pairwise PERMANOVA comparisons found prokaryote July and May communities to be more similar to each other than July and December (R^2^ of 0.04 and 0.14, respectively, *p* < 0.001). Cropping system, crop, N application and block all had minor yet significant effects (R^2^ = 0.06, 0.01, 0.05 and 0.02, respectively). Both time and cropping system affected fungal communities (R^2^ = 0.10 and 0.12, respectively, *p* < 0.001) with crop, N application and block also having relatively minor yet significant effects (R^2^ = 0.04, 0.01 and 0.01, respectively). The best separation could be observed as the CA system (blue) clustering in the top quadrats and the December communities (circles) clustering in the left-hand quadrats. Pairwise PERMANOVA also found July and May fungal communities to be more similar to each other than July and December (R^2^ of 0.04 and 0.11, respectively, *p* < 0.001). Finally, AMF were primarily affected by cropping system (R^2^ = 0.10, *p* < 0.001) more so than time (R^2^ = 0.02, *p* < 0.001). This was evident by the CON communities (black) separating from communities under the other three cropping systems. Crop, N application and block had little effect (R^2^ = 0.02, 0.01 and 0.02, respectively). Pairwise PERMANOVA comparisons of AMF communities also found July and May communities to be more similar than July and December (R^2^ of 0.01 and 0.02, respectively, *p* < 0.001). However, it should be noted that CLR-transformed data displayed non-homogenous variance in prokaryote, fungal and AMF multivariate matrices (Korkmaz *et al*., 2014) and consequently the reported *p* values, as with all *p* values, should be interpreted with caution (Colquhoun, 2014).

**Figure 4:**
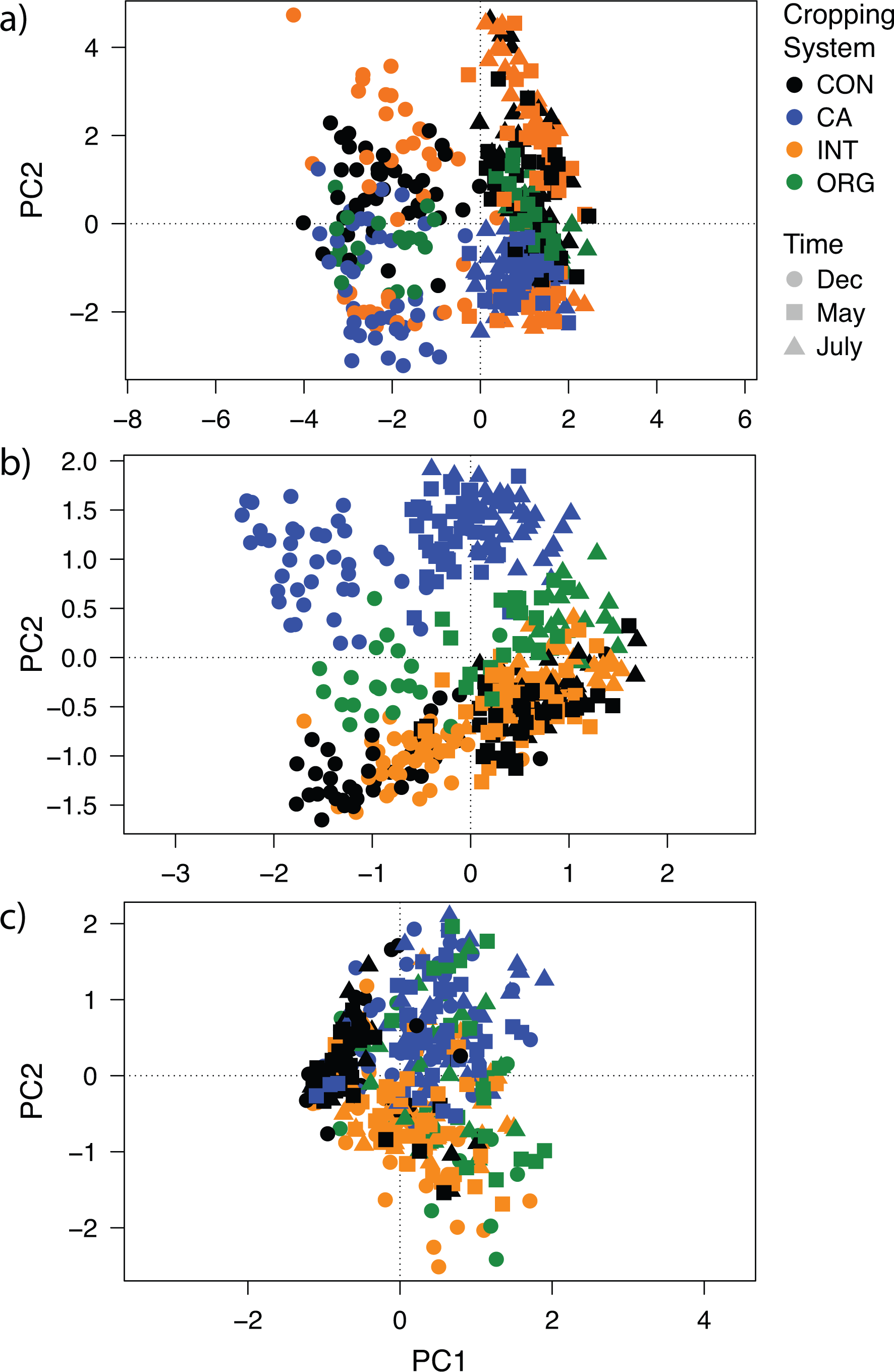
Ordinations of community composition for: a) prokaryotes, b) fungi and c) AMF.

### 3.2. Network analyses

Weighted networks of prokaryote, fungal and AMF communities were performed to visualise the stability of relationships between individual taxa under the four cropping systems, over time. Each node represents an OTU and each edge represents significant covariance. Table 3 lists node and edge numbers, mean and standard deviation of edge betweenness scores, clustering coefficients and the diameter of each network. Figure 5a are prokaryote networks for CON, CA, INT and ORG, respectively. While CON had the highest nodes, CA and INT had greater edges. Edge betweenness varied between the four systems. In the prokaryote networks, the clustering coefficients were relatively high under all cropping systems (0.51 – 0.57) due to the formation of two distinct, strongly covarying clusters. Notably, one cluster tended to be dominated by Actinobacteria and Alphaproteobacteria (brown and blue nodes, respectively), whilst the other cluster tended to host a diversity of Acidobacteria, Bacteroidetes, Verrucomicrobia and other phyla. In fungal networks, CON had lower nodes than CA and INT yet higher edges. The edge betweenness was consistently low in CON, however, which indicated edge lengths between CA, INT and ORG nodes tended to be shorter, and therefore that taxa covaried more strongly. Fungal clustering coefficients were all low, relative to prokaryotes, due to their networks tending toward a singular Ascomycota-dominated cluster with all other nodes sparsely connected and radiating outward (Figure 5b). Finally, the AMF CON network was most distinct from CA, INT and ORG with very low edges, high edge betweenness and similar clustering coefficient, despite having similar numbers of nodes. Visually this resulted in the AMF CON network having several densely packed, unconnected clusters including six nodes separate from the primary network. This differed markedly from the highly connected AMF networks under the other three systems, particularly the dense CA network.

**Figure 5:**
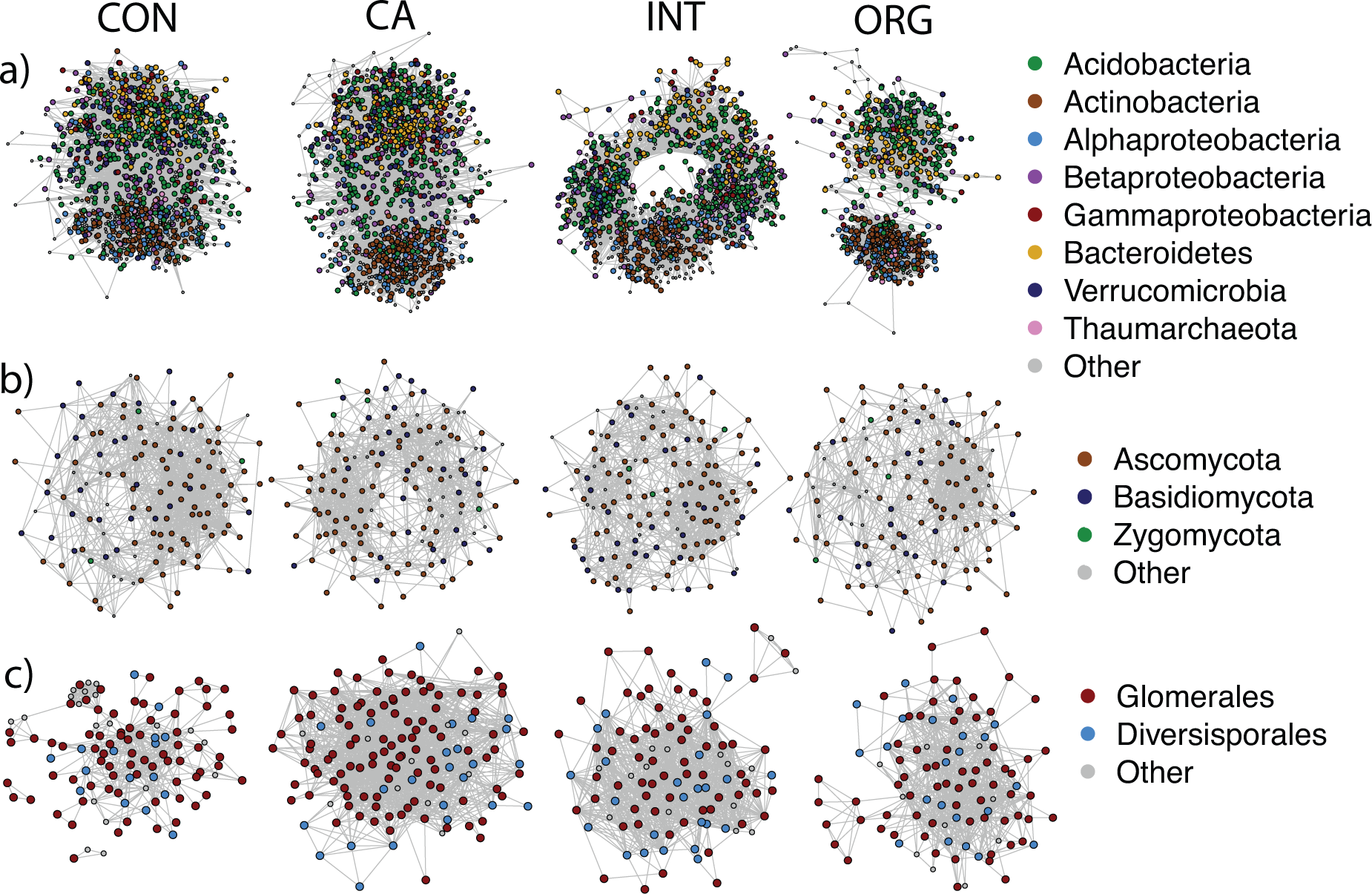
Weighted networks under CON, CA, INT and ORG for a) prokaryotes, b) fungi and c) AMF. Nodes of major taxonomic groups have been coloured and enlarged.

**Table 3:** Network properties separated by cropping system.

|  |  | CON | CA | INT | ORG |
| --- | --- | --- | --- | --- | --- |
| Prokaryotes | Nodes | 1257 | 1215 | 1231 | 706 |
|  | Edges | 28131 | 28317 | 30441 | 10926 |
| | Betweenness | 1619 ± 4022 | 1659 ± 4229 | 1588 ± 3785 | 1284 ± 4629 |
|  | Clustering coef. | 0.55 | 0.57 | 0.53 | 0.51 |
|  | Diameter | 5407 | 5024 | 5945 | 3180 |
| Fungi | Nodes | 133 | 140 | 142 | 125 |
|  | Edges | 1130 | 925 | 1017 | 732 |
| | Betweenness | 92 ± 101 | 104 ± 123 | 107 ± 117 | 100 ± 111 |
|  | Clustering coef. | 0.49 | 0.42 | 0.47 | 0.47 |
|  | Diameter | 1363 | 1241 | 1620 | 421 |
| AMF | Nodes | 109 | 139 | 105 | 100 |
|  | Edges | 407 | 1404 | 985 | 684 |
| | Betweenness | 100 ± 197 | 88 ± 95 | 70 ± 101 | 71 ± 108 |
|  | Clustering coef. | 0.45 | 0.41 | 0.51 | 0.44 |
|  | Diameter | 256 | 282 | 363 | 157 |

### 3.3. Stability of communities over time

Figure 6a, b and c show JS indices for prokaryote, fungal and AMF communities over time. Table 4 shows mixed effects model results of JS differences between cropping systems at December and May timepoints. The intercepts for each mixed effects model followed the trend prokaryotes > fungi > AMF. Within each group, July (harvest) was more similar to May (during growing season) than December (overwinter). For prokaryotes, only CA had a weakly positive effect on community similarity (0.57 and 0.62, respectively) relative to CON (0.55 and 0.59, respectively). However, community similarity was strongly improved in fungal communities under CA (0.49) and INT (0.47) relative to CON (0.44) during overwinter, with this positive effect lasting under CA (0.51) relative to CON (0.49) during the growing season. The most marked differences between cropping systems were related to the AMF communities, despite having the lowest JS values and therefore having the most dissimilar compositions at overwinter and during the growing season. Similarity was greatest under CA (0.22) and ORG (0.26) relative to CON (0.15) during overwinter, which further improved in CA, INT and ORG (0.36, 0.35, 0.32 respectively) relative to CON (0.24) during the growing season. Random effects only explained minor variation in JS (< 10%) with the exception of crop (wheat versus rapeseed (alfalfa for ORG)) playing a larger role in background variation for AMF resistance (22.1%).

**Figure 6:**
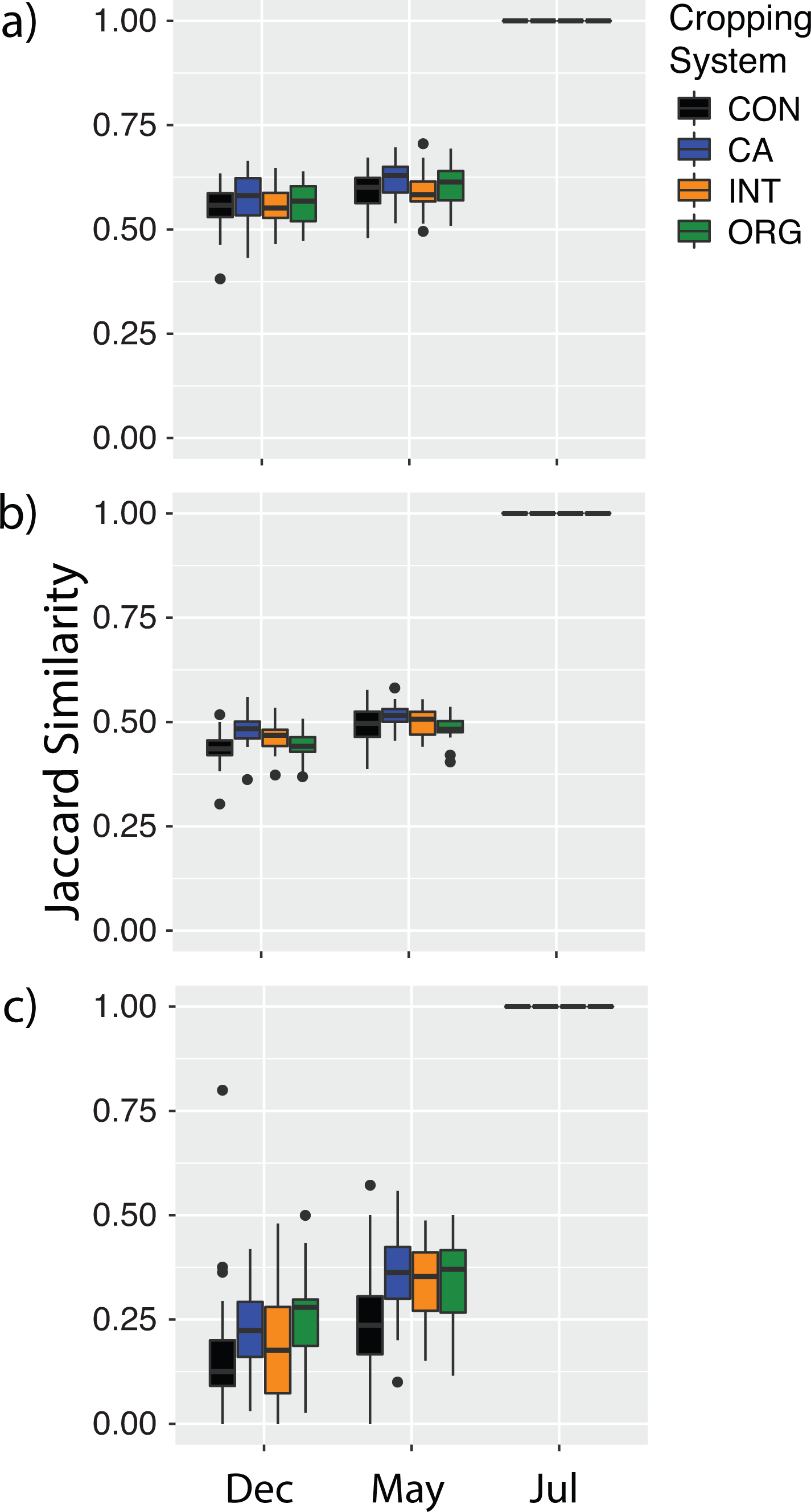
Observed JS over time and between cropping system for: a) prokaryotes, b) fungi and c) AMF.

**Table 4:**
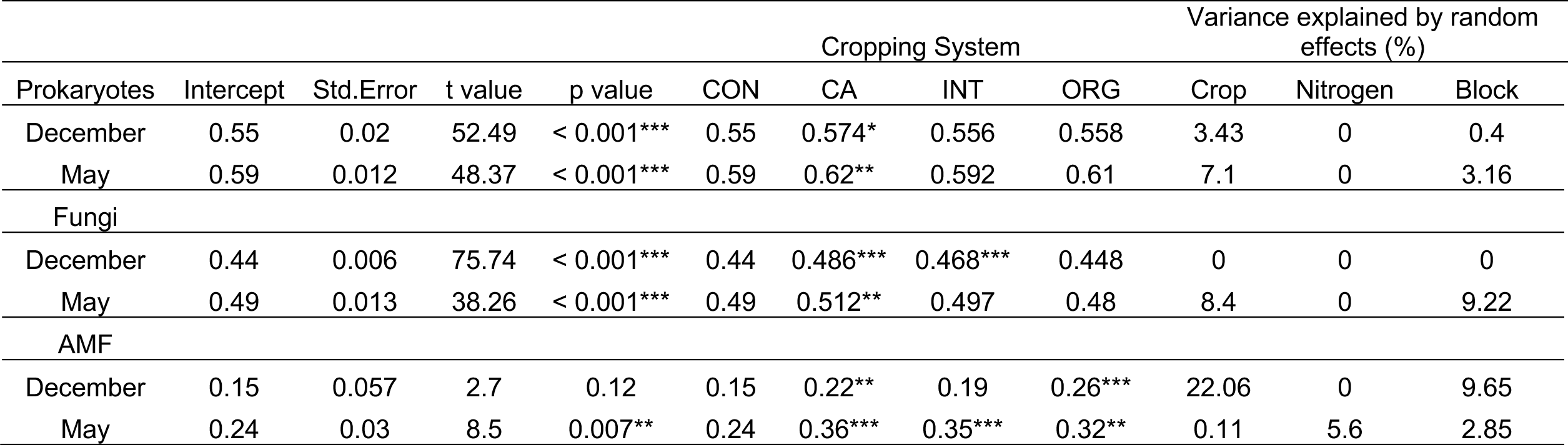
Mixed effects models of cropping system effects on JS at time point, with statistical significance noted as: (***) *p* < 0.001, (**) *p* = 0.001, (*) *p* < 0.05. Variance explained by random effects crop, year of N application and block are also shown.

## 4. Discussion

### 4.1. General trends in high taxonomic ranks across time and cropping system

The removal of a crop at harvest is likely to act as a disturbance event for soil communities in agricultural systems. We sought to investigate whether cropping systems that increased microbial alpha-diversity would also improve the stability of community composition over a growing season. The four cropping systems compared here were diverse: CON with frequent tillage, frequent pesticide and mineral N application; CA with no tillage, infrequent pesticide application, an alfalfa and red fescue cover crop, and both mineral N and legume rotation for fertilisation; INT with reduced tillage, infrequent pesticide and reduced mineral N application; and ORG with frequent tillage, no pesticides and a legume rotation for fertilisation.

In general, the dominant prokaryotic phyla in these soils reflected what is known of abundant phyla in agricultural systems. Individual phyla contain organisms representing considerable metabolic and physiological diversity, but conserved traits are observed (Finn *et al*., 2021) with trends in changing abundance and diversity detected at this high taxonomic rank in soil systems. Nevertheless, it should be recognized that the behaviour of specific taxa may deviate from these general trends. Bacteroidetes and Proteobacteria are dominant phyla in the wheat and rapeseed rhizosphere (Ai *et al*., 2015, Gkarmiri *et al*., 2017). Unsurprisingly, Bacteroidetes, and many Proteobacteria decreased markedly in December in the absence of a mature plant host. These phyla include copiotrophic taxa that are enriched by amino acids and monosaccharides (Fierer *et al*., 2007, Goldfarb *et al*., 2011, Ho *et al*., 2017) which are common root exudates that tend to be less available in bulk soils (Dennis *et al*., 2010, Kuzyakov & Blagodatskaya, 2015). Conversely, the Actinobacteria thrived in December (Figure 2). While this phylum is also present in both the wheat and rapeseed rhizosphere (Ai *et al*., 2015, Gkarmiri *et al*., 2017) it includes many saprotrophs that excel at degrading and consuming dead microbial and plant biomass (Barka *et al*., 2015). Thus, while the removal of the mature plant host appeared detrimental to phyla that included many plant-microbe symbionts, it was of benefit to saprotrophs. Furthermore, many Actinobacteria are well characterised psychrotolerant members of agricultural soils (Subramanian *et al*., 2016) and cold deserts (Siddiqui *et al*., 2013) and the traits that allow these organisms to tolerate low temperatures may have contributed to their predominance in over-wintering December soils.

The Thaumarchaeota, Planctomycetes and Acidobacteria were all affected by cropping system. These three phyla are notably enriched under low-intensity practices (Wessen *et al*., 2010, Ding *et al*., 2016, Li *et al*., 2017) while simultaneously decreasing under practices that involve adding mineral nitrogen (Fierer *et al*., 2012, Ramirez *et al*., 2012). Here, the relative abundance of Acidobacteria was greatest under ORG (25.9%) relative to the other systems (22.7, 21.4 and 22% for CON, CA and INT, respectively). The Acidobacteria are also present in the wheat and rapeseed rhizosphere but may not necessarily grow directly on plant root exudates, as shown through isotope tracing (Ai *et al*., 2015, Gkarmiri *et al*., 2017). It was hypothesised that the Acidobacteria are specialised to utilise recalcitrant plant organic matter in the rhizosphere, such as sloughed root cells. The capacity to utilise relatively recalcitrant plant matter would allow this taxonomic group to subsist through December until a new rhizosphere begins to take shape during the next growing season, and may explain why Acidobacteria were unaffected over the time of the growing season.

The most abundant fungal phylum was the Ascomycota (Figure 2). This phylum frequently dominates cropping and pasture systems (de Castro *et al*., 2008, Klaubauf *et al*., 2010, Ma *et al*., 2013). They are a highly diverse phylum (Lutzoni *et al*., 2004) that includes saprotrophs (Voriskova & Baldrian, 2013), symbiotic endophytes of wheat and rapeseed (Abdellatif *et al*., 2009, Behie & Bidochka, 2014, Gkarmiri *et al*., 2017), and a number of pathogens of wheat and other crops, including the genera *Phaesophaeria*, *Alternaria* and *Fusarium* (Berbee, 2001). Here they were particularly enriched in the CA, INT and ORG systems (58.5, 58.8 and 61.6%, respectively) relative to CON (54.5%). The second most abundant fungal phyla here were the Zygomycota. This differs from other agricultural systems that show Basidiomycota as the second most dominant group (de Castro *et al*., 2008, Klaubauf *et al*., 2010). The Zygomycota have been noted as dominant saprotrophs when plant organic matter is depleted of lignin relative to cellulose, toward the latter stages of decomposition (Osono & Takeda, 2001, Vivelo & Bhatnagar, 2019). Here, the effect of time on Zygomycota favoured an increase in their abundance in May and July where a growing plant may have supplied them with lignin-depleted plant material.

Finally, analysis of AMF Glomeromycota identified Glomerales and Diversisporales to be the dominant orders. The overwhelming dominance of Glomerales in agricultural soils has been noted previously where three geographically separate soils under three different crops were dominated by > 90% Glomerales (Helgason *et al*., 1998). In arid agricultural soils the community can reflect roughly 50% Glomerales / 50% Diversisporales (Li *et al*., 2007) and abiotic stress (as fungicides, pollutants) also drive communities to be dominated by Glomerales and Diversisporales (Lenoir *et al*., 2016). However, it should be noted that certain studies have found Glomerales and Paraglomerales to dominate agricultural systems (Sale *et al*., 2015, Banerjee *et al*., 2019). Interestingly, brief ‘blooms’ of Diversisporales were apparent in December and several ORG May and July samples (Figure 2). These shifts may be a technical artefact arising from slight variation in relative abundance (%) being more obvious within the species poor (∼ 250 OTUs) AMF community.

### 4.2. Time and cropping system effects on alpha-diversity and community composition

We first sought to investigate whether certain cropping systems would be beneficial for microbial alpha-diversity. Only time, and not cropping system, affected prokaryote alpha-diversity, with low biodiversity in December during the combined stressors of an absence of a mature plant host and low ambient temperature. Conversely, both richness and Shannon were highest in July near the culmination of harvesting with a mature plant host and mesophilic temperatures (Figure 3a). Although this study did not separately sequence the rhizosphere and bulk soil microenvironments, the presence of a mature plant likely increased the alpha-diversity, which further increased from May to July prior to harvest. Time was also the greatest determiner of prokaryote community composition, with December communities distinct from May and July

(Figure 4a). It is unclear whether prokaryote biodiversity is generally greater in the rhizosphere than bulk soil (Berg & Smalla, 2009, Philippot *et al*., 2013) with some studies suggesting selection for specific rhizosphere taxa reduces overall biodiversity (Ai *et al*., 2015) and others showing enriched biodiversity relative to bulk soil (McPherson *et al*., 2018). Here, the presence of a mature crop was likely beneficial to prokaryote alpha-diversity and in shaping the community composition.

While fungal alpha-diversity was high in both December and July, cropping system played a greater role in fungal richness and Shannon index than observed for prokaryotes (Figure 3b). Specifically, CA and INT were beneficial for richness, while ORG was detrimental for diversity (Table 2). The ORG practice had significantly lower TOC than the other practices and produced the lowest crop biomass t ha^-1^ year^-1^. Conversely, the TOC-rich CA practice had the strongest positive effect on fungal richness and resulted in a unique community composition (Figure 4b). As soil TOC is predominantly plant C (Dungait *et al*., 2012) and assuming the majority of the Ascomycota, Basidiomycota and Zygomycota are saprotrophs reliant on plant debris from crops, the low TOC may be responsible for the decreased Shannon observed under ORG. At the global scale, climate and plant diversity are the most important drivers of fungal biodiversity (Tedersoo *et al*., 2014). However, at more local scales, for example within northern China and within the Mediterranean, TOC is the most important driver of fungal biodiversity (Persiani *et al*., 2008, Liu *et al*., 2015). The abundance of Ascomycota and Zygomycota phyla were also shown to be particularly dependent on TOC (Liu *et al*., 2015). Thus, the TOC promoted by the differential cropping systems likely influenced overall fungal biodiversity.

The alpha-diversity of AMF followed clear trends of increasing from December, through May to July prior to harvest, with the CON system demonstrating consistently poor richness and Shannon indices. These effects were much stronger on the AMF than the other taxonomic groups (Table 2). Furthermore, community composition under the CON practice was most dissimilar to the other practices (Figure 4c). Tillage, and frequent pesticide application in the case of CON, have the potential to severely limit the alpha-diversity of AMF (Sale *et al*., 2015, Gottshall *et al*., 2017, Banerjee *et al*., 2019). Conversely, the CA and ORG systems promoted AMF alpha-diversity the most, likely as a consequence of incorporating a cover crop (Vukicevich *et al*., 2016, Hontoria *et al*., 2019) and reduced management intensity of mineral N fertilisation and pesticide application (Banerjee *et al*., 2019). The results here are in strong agreement with previous research on positive and negative drivers of AMF communities.

Therefore, cropping system positively affected fungal richness, with CON being consistently relatively low and CA relatively high, while fungal Shannon diversity was low under ORG relative to CON. The AMF alpha-diversity increased under all systems relative to CON.

### 4.3. Cropping system effects on relationships between taxa

Another community aspect investigated was whether cropping systems could promote more stable relationships between individual taxa, which can be visualised through network analyses. Interestingly, all prokaryote networks showed the formation of two distinct clusters of significantly co-varying taxa (Figure 5a) accompanied by high clustering coefficients. Regardless of cropping system, one cluster tended toward low biodiversity dominated by potentially saprotrophic Actinobacteria and Alphaproteobacteria (the December overwinter cluster), while the other cluster tended toward higher biodiversity with all Proteobacteria classes, Bacteroidetes and Acidobacteria (the crop-associated July and May cluster). Taken together, these analyses suggest that while the absence of a mature plant and psychrotolerance potentially decreased the overall alpha-diversity in December, the prokaryote communities were flexible in that a novel community was established. Then, in the presence of a mature plant, a distinct crop-associated community emerged with increased alpha-diversity until harvest in July. It is likely that other factors associated with seasonal changes also contributed as community drivers (e.g. soil moisture).

Unlike the prokaryotes, the fungal community did not form distinct clusters of significantly covarying taxa – rather a central core of Ascomycota taxa was consistently present with other less-densely connected taxa radiating outward toward the periphery (Figure 5b). Despite being the second most dominant phylum, only few individual, scattered Zygomycota nodes were in the networks, indicating that only few of these taxa covaried significantly with others across time. As discussed above, the low TOC under ORG, which likely resulted in low biodiversity, was also likely responsible for sparse network connectivity between taxa in the ORG network, and therefore yielded fewer potential stable relationships between taxa.

Recently, Banerjee *et al*., (2019) found a negative relationship between increasingly intensive agricultural management practices and node, edge and edge betweenness values within AMF community networks. Here the high-intensity CON system yielded the most distinct AMF network notable for its sparse connectivity. In contrast, the relatively low-intensity, permanent cover crop CA resulted in the most connected CA networks, indicative of strong and stable relationships between AMF over time.

Therefore, the strongest relationships between taxa were only observed for prokaryotes, and this was driven by temporal changes between overwinter *versus* crop-associated communities. An effect of cropping system on network structure was primarily demonstrated by the AMF CON and CA networks, which were either sparsely or highly connected, respectively.

### 4.4. Stability of soil microbial communities over time

The factors that confer stability to soil microbial communities have received a great deal of attention due to the role of microbes as drivers of ecosystem processes (Griffiths and Philippot, 2012, and references therein). Disturbance can come in many forms – drought (Manzoni *et al*., 2012), fire (Banning & Murphy, 2008), metal (Philippot *et al*., 2008) and organic pollutants (Girvan *et al*., 2005). Increasing biodiversity of a community can strengthen its capacity to respond to a disturbance event, termed the ‘insurance hypothesis’ (Ives *et al*., 2000). Here we considered crop harvesting followed by a fallow period as a disturbance, and aimed to determine whether cropping systems that promoted alpha-diversity would also demonstrate higher similarity in composition over a growing season, *i.e.* they would be more stable. This effect on community composition was measured with JS as a robust and simple pairwise metric of community similarity (Scheiner, 1992) although it should be noted that it is unweighted by relative abundance. As one would expect, in all three groups, the similarity to the harvest community increased over time as communities transitioned from overwinter. Another general trend was that the composition of prokaryotes was most similar (*i.e.* stable) over time, fungi intermediate and AMF relatively poor (Figure 6, Intercepts in Table 4). This agreed with the insurance hypothesis at a broad taxonomic level, with the species-rich prokaryotes demonstrating greater stability than species-poor AMF, regardless of cropping system.

There was a positive benefit to stability to all groups under CA, with this effect relatively more pronounced in the fungi and AMF. Soil aggregation and TOC can provide isolated microhabitats and/or support growth of microorganisms post-disturbance, and these edaphic properties have been strongly linked to stability (Griffiths *et al*., 2005, Zhang *et al*., 2010). As soils under CA were no-till and had a permanent cover crop, and likely had better soil aggregation than annually tilled CON and ORG (West & Post, 2002), this may explain the generally higher stability observed under this cropping system. In contrast, the combination of annual tillage, mineral N fertilisation and frequent pesticide application was likely responsible for the poor stability of AMF observed under CON. Finally, it is possible that the stability of all communities measured here are high relative to other forms of disturbance in soil systems if communities have adapted to annual crop harvesting and fallow periods (Griffiths & Philippot, 2013). With the exception of the cover crop in CA soils, each practice endures an annual fallow period which has been ongoing for 18 years. This may act as a form of pressure to strictly select for microbial species that can both survive during this fallow period in the absence of an actively growing crop and respond rapidly to a new crop during the growing season. Thus, one could reasonably suppose that the communities here are somewhat adapted to recovering after crop removal. Even so, crop removal and the ensuing overwinter period had dramatic effects on the composition of all communities, relative to the harvest period, particularly for the AMF.

In summary, relative to CON, CA had a consistently positive effect on prokaryote and fungal community stability over time. The AMF demonstrated greater stability under all cropping systems relative to CON. Thus, practices that increased alpha-diversity in fungi (and AMF) also appeared to convey more stable community compositions. The increased stability in prokaryotes under CA, without increased alpha-diversity, suggests that increasing alpha-diversity is not, however, absolutely necessary for a more stable community.

## 5. Conclusions

Presented here is an investigation of prokaryote, fungal and AMF soil communities that demonstrated markedly different changes over a growing season, under four agricultural cropping systems. These cropping systems had been implemented for 18 years, giving ample time to differentially shape these communities. Prokaryotes were flexible and formed a distinct saprotroph-dominated community in the absence of a mature crop overwinter, and a crop-associated community in the presence of a mature crop. This relatively species rich group demonstrated the highest stability over time.

The CA system was associated with the most diverse fungal community, which also increased its stability over the course of the growing season. Finally, the CON practice had a particularly detrimental effect on AMF alpha-diversity and stability relative to all other cropping systems. In conclusion, cropping systems that promoted alpha-diversity led to increased stability of fungal (and AMF) communities over the growing season, with prokaryotes being relatively insensitive to the effect of soil management.

## Acknowledgements

We would like to thank INRAE, Versailles-Grignon for use of the La Cage field trial, and Gilles Grandeau for assistance with soil sampling.

